# Transcriptional control by Sall4 in blastocysts facilitates lineage commitment of inner cell mass cells

**DOI:** 10.1101/194852

**Authors:** Anzy Miller, Sarah Gharbi, Charles Etienne-Dumeau, Ryuichi Nishinakamura, Brian Hendrich

## Abstract

The enhancer-binding zinc finger transcription factor Sall4 is essential for early mammalian postimplantation development and plays important roles in lineage commitment of embryonic stem cells. Enhancer binding by Sall4 results in transcriptional activation of some genes, but repression of others. Exactly how cells in preimplantation stage embryos use this transcriptional modulatory activity of Sall4 during early developmental transitions has not been determined. Using single cell gene expression analyses we show that Sall4 is required to maintain the gene regulatory network in inner cell mass (ICM) cells prior to lineage commitment. Although Sall4 is not required for ICM cells to adopt a correct epiblast or primitive endoderm gene expression profile, in the absence of Sall4 early ICM cells commit to either lineage at reduced frequency. We propose a model whereby Sall4 activity sets the stage for efficient progression from the uncommitted ICM progenitor state by modulating the gene regulatory network in early ICM cells.

## Introduction

Mammalian pre-implantation development is a paradigm of regulative self-organisation and differentiation. The first cell fate decision in mammalian development occurs when the outer cells of the 16-to-32 cell embryo become destined to form trophectoderm (TE), which will go on to form extraembryonic tissue, and the inside cells become the inner cell mass (ICM), which retain both embryonic and extraembryonic potential (Gardner et al., 1973; Gardner and Rossant, 1979; Papaioannou, 1982; Tarkowski and Wroblewska, 1967). At this point the embryo forms a blastocyst, with TE cells lining the outside of the structure and the ICM cells forming a mass at one end of the embryo. The second cell fate decision occurs in mid stage blastocysts as cells of the ICM form either the extraembryonic primitive endoderm or the embryonic epiblast cells, which retain pluripotency (Gardner and Rossant, 1979).

The transcriptional changes that occur during these first two cell fate decisions have recently been profiled using single cell gene expression techniques (recently reviewed in (Kumar et al., 2017)). The identification of transcription factors important for these decisions has provided some indication of how these transcriptional changes are controlled (Moris et al., 2016).

Sall4 is a C2H2-type zinc finger transcription factor which plays important roles in various aspects of mammalian development. *Sall4* is abundantly expressed in early embryos and germ cells, but in fully differentiated tissues there is little or no expression (Kohlhase et al., 2002; Tsubooka et al., 2009). Mutations in *Sall4* show haploinsufficiency, leading to anorectal and cardiac anomalies in mice (Koshiba-Takeuchi et al., 2006; Sakaki-Yumoto et al., 2006), and causing Okihiro/Duane-Radial Ray and IVIC syndromes in humans (Al-Baradie et al., 2002; Kohlhase et al., 2002; Sweetman and Munsterberg, 2006). *SALL4* expression is aberrantly activated in multiple tumour types, leading to speculation that it may play a facilitating role in tumour progression (Tatetsu et al., 2016; Zhang et al., 2015).

Sall4 predominantly associates with enhancer sequences in mouse embryonic stem (ES) cells (Miller et al., 2016; Xiong et al., 2016). Sall4 does not appear to recognise any specific DNA sequence motif, but rather is found at sequences bound by other enhancer-binding transcription factors in ES cells, such as Nanog, Oct4 (encoded by the *Pou5f1* gene), Klf4 and Esrrb. Sall4 can stimulate expression of some pluripotency-associated genes in mouse ES cells, but it is not necessary for the maintenance of the pluripotency gene regulatory network (GRN) (Miller et al., 2016; Wu et al., 2006; Zhang et al., 2006). Rather, Sall4 acts to prevent inappropriate activation of the neural-specific GRN in undifferentiated ES cells, and is also necessary for appropriate activation of gene expression patterns in differentiating ES cells (Miller et al., 2016).

Sall4 plays an essential role in early mouse development. *Sall4*-null blastocysts are produced in normal ratios from heterozygous intercrosses, appear morphologically normal, and are able to implant into the uterus or to attach to substrate in vitro (Elling et al., 2006; Sakaki-Yumoto et al., 2006). Despite this, these embryos undergo a dramatic developmental failure at the periimplantation stage, such that little or no embryo-derived tissue remains at early post-implantation stages (Elling et al., 2006; Sakaki-Yumoto et al., 2006). Maternal *Sall4* protein and transcript are detectable in zygotes, but no maternally-derived protein or mRNA is detectable in cleavage stage embryos (Xu et al., 2017; Zhang et al., 2006). Zygotic Sall4 protein becomes detectable in 8-cell embryos and remains expressed through the rest of preimplantation development (Elling et al., 2006; Sakaki-Yumoto et al., 2006; Xu et al., 2017; Zhang et al., 2006), but it is not known what role is played by Sall4 in pre- and peri-implantation embryos.

In this study we investigated Sall4 function in mouse preimplantation development. Using single cell gene expression analysis of key genes associated with the first cell fate decisions, we find that Sall4 is required for appropriate gene expression in ICM cells of early blastocysts. Though cells are able to partially compensate for this gene expression defect in late stage blastocysts, mutant embryos show a reduced efficiency of lineage commitment as cells attempt to undergo the second lineage decision, that of epiblast or primitive endoderm. Blastocysts are able to make both epiblast and primitive endoderm cells in the absence of Sall4, but they do so with a reduced frequency. Thus transcriptional control by an enhancer binding protein in unspecified ICM cells primes these cells for efficient subsequent lineage commitment.

## Results and Discussion

### Sall4 is required for appropriate gene expression in early blastocysts

Maternally derived Sall4 protein was undetectable in homozygous null embryos at the 16-cell stage (Fig. 1A) as reported (Elling et al., 2006; Sakaki-Yumoto et al., 2006). *Sall4*-null early blastocysts (embryonic day 3.5; E3.5) appeared morphologically normal, displaying ubiquitous Oct4 expression and normal cell numbers (Fig. 1B, C). This indicated that zygotic Sall4 is not required for proliferation, cell viability, or normal Oct4 protein distribution in early blastocysts. At the morula and early blastocyst stages *Sall4*-null embryos contained significantly fewer Nanog-expressing cells than did wild type or heterozygous embryos, though these cells expressed Nanog at normal levels (Fig. 1B, C, S1A, B, C). By the mid-blastocyst stage (E3.75), both wild type and mutant embryos had restricted Nanog expression to the ICM and Cdx2 expression to outer cells (Fig. 1D), consistent with successful completion of the first cell fate decision, that of trophectoderm vs ICM. At this point wild type and *Sall4*-null embryos expressed Nanog in a similar proportion of cells (~30%, Fig. 1D, E, S1D). Notably, *Sall4*-mutant mid-stage blastocysts failed to activate the primitive endoderm marker Gata4 (Fig. 1D, E). Sall4 is therefore dispensable for the first cell fate decision, but its absence correlates with a reduced number of Nanog-expressing cells in early blastocysts, and of Gata4-expressing cells in mid-blastocysts.

**Figure 1.**
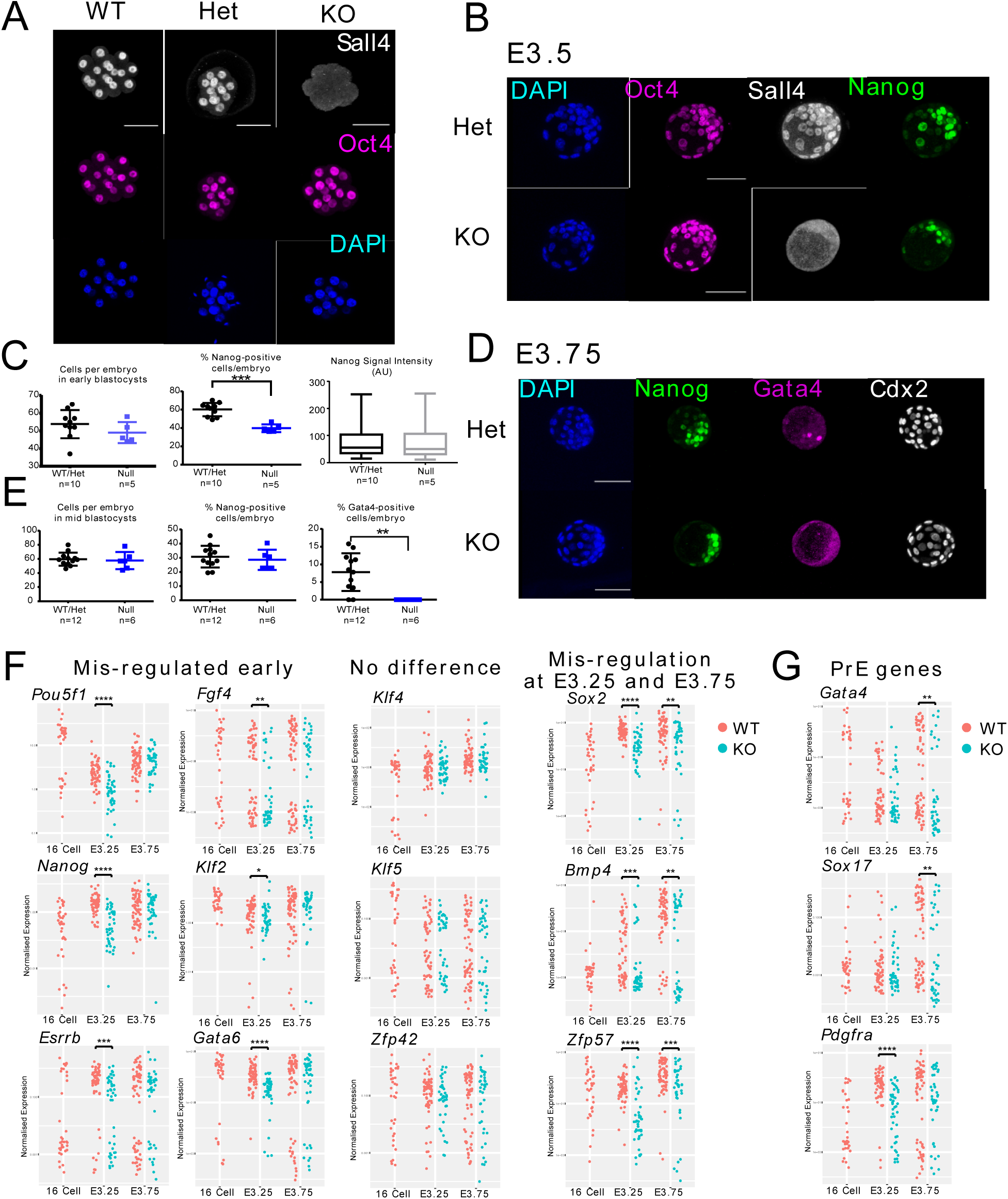
Altered gene expression in *Sall4*-null blastocysts. (A)Representative projection images of anti-Sall4 (white), anti-Oct4 (magenta) and DAPI (blue) staining in WT (n=3), Sall4^Δ/+^ (Het, n=2) and Sall4^Δ/Δ^ (KO, n=3) 16 cell stage embryos. Scale bars = 50μM. (B)Representative projection images of DAPI (blue), anti Nanog (green), anti-Oct4 (magenta) and anti-Sall4 (white) staining in control (Het; N = 10) and Sall4^Δ/Δ^ (KO; N = 5) early blastocyst embryos (E3.5). Scale bars = 50μM. (C)Quantification of the number of cells per embryos from (B) (left), the percentage of Nanog positive cells per embryo (middle), and intensity values for the Nanog positive cells from (B) for control (Wt/Het) and KO embryos (right). Significance was determined using a Mann-Whitney test for all three comparisons (*** P≤ 0.001). (D)Representative projection images of anti-Nanog (green), anti-Gata4 (magenta), anti-Cdx2 (white) staining, plus DAPI (blue) in control (Het; n=7) and Sall4^Δ/Δ^ (KO, n=6) E3.75 embryos. Scale bar = 50μM. (E)Quantification of the number of cells per embryos from (D) (left), the percentage of Nanog positive cells per embryo (middle), and percentage of Gata4 positive cells per embryo (right) for control (Wt/Het) and KO embryos. Significance was determined using a Mann-Whitney test (** P≤ 0.01). (F)Single cell gene expression analysis from E3.25 ICM WT/Het and KO embryos at indicated stages, with Log^10^ expression normalised to that of housekeeping genes (see Methods) on the y-axis. Each dot represents a single cell, and each plot shows data for the indicated gene. Statistical significance was calculated using a Kruskall-Wallis test (Supplemental Table 1), * indicates P ≤0.05, ** P ≤0.01, *** P ≤0.001, **** P <=0.0001. Published data from wild type 16-cell embryos are included for comparison (O’Shaughnessy-Kirwan et al., 2015). (G)Same as in (F), but for genes expressed in primitive endoderm cells.

Sall4 is an enhancer-binding transcription factor, so we suspected that the altered protein expression patterns observed in *Sall4*-null embryos would be indicative of altered transcription in early embryonic cells. To test this we measured expression of genes associated with early embryonic lineages by quantitative reverse-transcription PCR (qRT-PCR) in individual ICM cells isolated from early and mid-stage blastocysts. Overall, the transcript levels of 32 genes were measured in 130 control (wild-type or het) cells and 85 *Sall4*-null cells. Expression data from wild type 8-and 16-cell embryos produced through identical methods (O’Shaughnessy-Kirwan et al., 2015) was included in the analyses to provide a developmental reference point.

*Sall4*-null cells from early stage ICMs (E3.25) showed reduced expression of many genes associated with the pluripotency GRN including *Nanog, Pou5f1, Sox2* and *Esrrb*, but not *Klf4, Klf5* or *Zfp42* (Fig. 1F). The expression levels of some genes, including *Pou5f1, Nanog, Sox2*, and *Gata6*, were reduced in *Sall4*-null cells compared to wild type cells. Other genes, such as *Esrrb, Bmp4* and *Zfp57*, were expressed in a reduced proportion of cells in mutant embryos (Fig. 1F). Notably, the reduced transcription of *Nanog* and *Pou5f1* detectable by qRT-PCR did not result in adecrease in protein levels detectable by immunofluorescence (Fig. 1B, C). Gene expression defects were less pronounced in mid-stage blastocysts (E3.75), with *Nanog, Pou5f1, Gata6* and *Klf2* all showing restoration of normal expression levels at the later stage (Fig 1F). Nevertheless, ICM cells from mid-stage *Sall4*-null blastocysts continued to express reduced levels of *Sox2, Bmp4*, and *Zfp57*, and showed a defect in activation of the primitive endoderm markers *Gata4* and *Sox17* (Fig. 1F). Importantly, the expression patterns seen in *Sall4*-null cells at the early blastocyst stage did not resemble that of either later or earlier wild type embryos, indicating that *Sall4*-null ICM cells are not simply developmentally advanced nor delayed, but rather display widespread alterations in gene expression.

### Impaired primitive endoderm formation in *Sall4*-null embryos

Early ICM cells express both Nanog and Gata6, but between the 32 and 120-cell stage cells asynchronously and irreversibly adopt the fate of either epiblast or primitive endoderm and silence Gata6 or Nanog, respectively (Chazaud et al., 2006; Grabarek et al., 2012; Nichols et al., 2009; Plusa et al., 2008; Saiz et al., 2016; Schrode et al., 2014). Mid-stage (E3.75) ICM cells appear to segregate into Nanog- and Gata6-expressing populations normally in the absence of Sall4 (Fig. 2A), and re-establish normal levels of *Nanog, Gata6* and *Pdgfra* transcripts (Fig. 1F, G), indicating Sall4 is dispensable for this early stage of the second cell fate decision. Yet in the absence of Sall4 ICM cells showed impaired activation of primitive endoderm genes *Gata4* and *Sox17*, and did not express Gata4 protein (Fig. 1D, E, G), likely indicating a defect in successful establishment of the primitive endoderm lineage.

**Figure 2.**
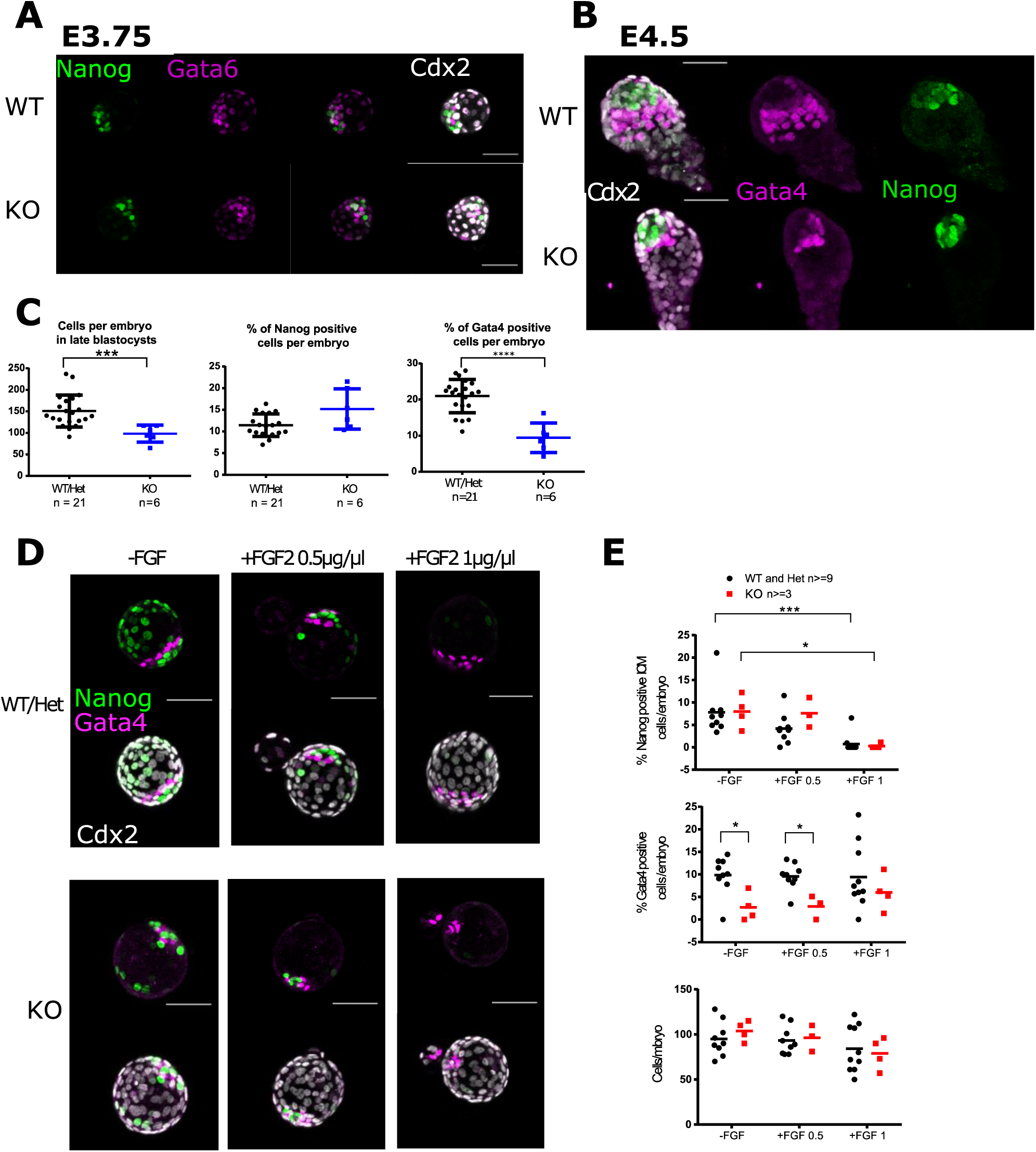
Sall4 facilitates primitive endoderm formation in implanting blastocysts. (A)Representative projection images of anti-Nanog (green), anti-Gata6 (magenta) staining, along with composite images of anti-Nanog and anti-Gata6 together, plus the inclusion of anti-Cdx2 (white) staining in control embryos (WT n=4) and Sall4^Δ/Δ^ (KO, n=5) E3.75 embryos. Scale bar = 50μM. (B)Representative projection images of anti-Nanog (green), anti-Gata4 (magenta), anti-Cdx2 (white) staining in control (WT n=9) and Sall4^Δ/Δ^ (KO, n=6) E4.5 late blastocysts. Scale bar = 50μM. (C)Quantification of the number of cells per embryo (left); the percentage of Nanog positive cells per embryo (middle); and the percentage of Gata4 positive cells per embryo at 4.5 dpc. *** P ≤0.001, **** P ≤0.0001 by Mann-Whitney test. (D)Representative confocal projection images of control (WT or Het) and Sall4^Δ/ Δ^ (KO) embryos flushed at 2.5dpc and cultured for 48hours in media only or with 0.5 μg/μl or 1 μg/μl Fgf2. Embryos were stained with anti-Gata4 (magenta), anti-Nanog (green) and anti-Cdx2 (white). Scale bar = 50μM. N=9 for WT or Het embryos cultured without Fgf2 and in 0.5μg/ml Fgf2, n=10 for those in 1μg/ml Fgf2. N=4 for KO embryos cultured without Fgf2 and in 1μg/ml and n=3 for those cultured in 0.5μg/ml Fgf2. (E)Percentage of Nanog positive (top) or Gata4 positive (middle) cells per embryo for control (WT and Het) or Sall4-null (KO) embryos after culture in indicated conditions for 48 hours. Each point represents one embryo. Statistical significance was tested using an unpaired Mann-Whitney test: * P ≤0.05, *** P ≤0.001.

By the late blastocyst stage (E4.5) *Sall4*-null embryos activated expression of primitive endoderm makers in a subset of cells, but still contained significantly fewer Gata4-expressing primitive endoderm cells than did wild type embryos, and those Gata4/Gata6/Sox17-expressing cells present did not appear to form an organised cell layer (Fig. 2B, C, S1F). Mutant embryos were smaller than wild type or heterozygous embryos and displayed an increased number of apoptotic cells, but contained a normal proportion of Nanog-expressing, presumptive epiblast cells per embryo (Fig. 2 B, C, S1E). Sall4 is thus dispensable for appropriate tissue segregation in peri-implantation stage embryos, but in its absence a reduced proportion of primitive endoderm cells is produced, thus skewing the epiblast to primitive endoderm ratio from 1:2 in wild type embryos to 1.5:1 in mutants.

Normal PrE formation is dependent upon FGF signalling, such that mutations in the Fgf signalling pathway lead to a complete loss of primitive endoderm formation during early development (Chazaud et al., 2006; Kang et al., 2017; Molotkov et al., 2017). Forced stimulation of Fgf signalling by culturing embryos in the presence of Fgf2 skews ICM cell fate commitment away from the epiblast fate (Le Bin et al., 2014; Saiz et al., 2016; Yamanaka et al., 2010). To test whether Sall4 plays an important role in the response to Fgf signalling, 8-cell embryos were cultured for 48 hours in two different concentrations of Fgf2 (or without Fgf2 as a control), and cell specification was assessed by staining for lineage-specific markers (Fig. 2D). Culturing embryos in Fgf2 resulted in a reduction in Nanog-expressing, presumptive epiblast cells, while having no effect on the number of cells per embryos in any condition or genotype (Fig. 2D, E). We therefore find no evidence that Sall4 facilitates Fgf signalling responsiveness in ICM cells.

### Sall4-dependent control of gene expression in unspecified ICM cells facilitates subsequent lineage commitment

Sall4 is important for primitive endoderm formation in peri-implantation stage blastocysts, yet gene expression patterns in *Sall4* null embryos are most divergent from wild type at the early blastocyst stage (Fig. 1F). We hypothesised that Sall4-mediated transcriptional control in early blastocysts is important for these early cells to acquire competence to undergo primitive endoderm formation. To test this hypothesis we interrogated our single cell gene expression data to determine at what stage(s) wild type and mutant cells differ based upon gene expression. The single cell gene expression data can be visualised in a diffusion plot (Fig. 3A). Including expression data from cells obtained from 8- and 16-cell embryos (O’Shaughnessy-Kirwan et al., 2015) provides a clear developmental orientation, with the least differentiated cells clustering in the lower right hand corner of the plot (Fig. 3A). Developmental progression then proceeds up and to the left, whereupon the population splits and cells occupy one of two different trajectories, representing cells committed to the epiblast or primitive endoderm lineages. *Sall4*-null cells from E3.25 ICMs (purple) occupy a more primitive position than do wild type cells, in that most cluster closer to the bottom right corner than do any of the wild type cells (Fig. 3A). The majority of cells from E3.75 *Sall4*-null ICMs (pink) occupy a position similar to that occupied by the E3.25 wildtype cells. Unlike early wild-type cells, a proportion of E3.75 *Sall4*-null cells occupy positions indicative of having committed to epiblast or primitive endoderm, similar to that of their wildtype counterparts.

**Figure 3.**
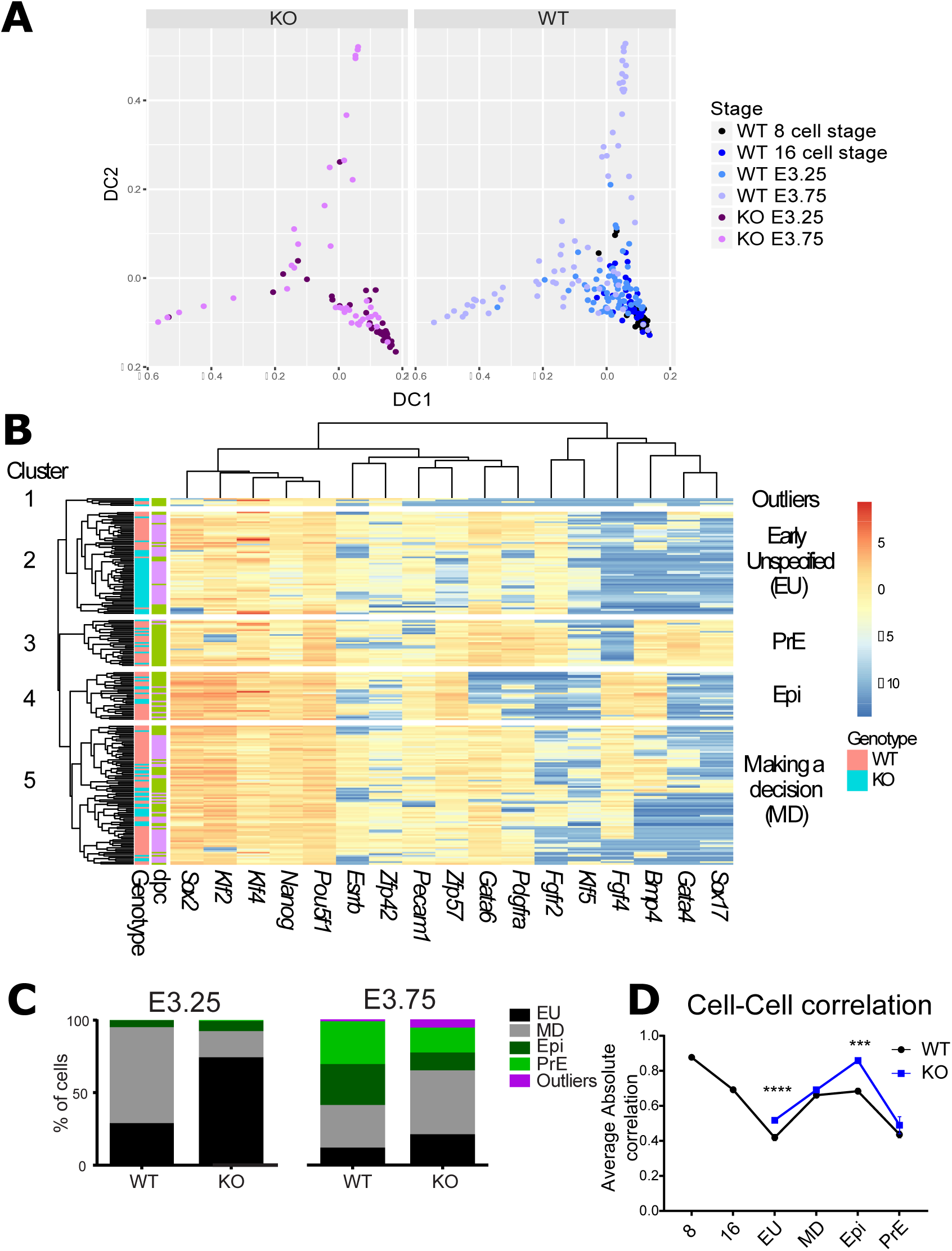
Single cell gene expression analysis of wild type and Sall4 null ICM cells. (A)Diffusion plot of single cell gene expression data for Sall4-null (KO, left) and wild type (WT, right) ICM cells. Points represent individual cells of the indicated stages and genotypes. 8- and 16-cell data is taken from O’Shaughnessy-Kirwan et al., 2015. (B)Heat map constructed from single ICM cell expression data based on hierarchical clustering. Normalised Ct values refer to –ΔCt values normalised to housekeeping genes (*Ppia, Gapdh* and *Hprt*). Each row is one cell with the genotype and the time isolated indicated by the annotation panel on the left and key on the right (dpc = days post coitum). Each column shows expression of one gene. The cells cluster into 5 groups which have been labelled according to their expression profiles: Early unspecified (EU), Making a decision (MD), Epiblast (Epi), Primitive Endoderm (PrE) and the outliers. (C)The percentage of cells in each cluster from (B) at each time point (E3.25 or E3.75) for KO and control (WT, including cells from Het embryos) ICM cells. (D)Absolute average correlation (see Figure S3) between cells for each developmental stage is plotted for control (WT, black) and Sall4-null (KO, blue) cells. CV for 8- and 16-cell stages were calculated using data from O’Shaugnessy-Kirwan et al 2015. Error bars represent SEM, and significance was determined using one-way ANOVA followed by Sidak’s multiple comparison test. *** P≤0.001, **** P≤0.0001.

Unsupervised clustering of the expression data divided the cells into 5 major clusters (Fig.3B). Cluster 1 is an outlier group containing five cells from 3.75 dpc embryos showing very low expression levels of most genes. We suspect that these cells may have failed to properly specify any lineage and hence may be destined for apoptosis. Cluster 2 predominantly consists of cells from early blastocysts which express markers of naïve cells, but low levels of markers associated with mature lineages (e.g. *Bmp4, Zfp57*). We infer that this cluster represents <u>e</u>arly <u>u</u>nspecified (EU) cells. Clusters 3, 4 and 5 exist on a separate branch of the tree from the EU cells. Clusters 3 and 4 contain almost exclusively cells from late blastocysts and express high levels of primitive endoderm or epiblast markers, respectively. We infer that these clusters represent cells committed to the primitive endoderm (PrE-Cluster 3) or epiblast (Epi – Cluster 4) lineage, respectively. Cluster 5 contains a mix of early and late blastocyst cells, most of which have activated *Fgf4* and *Zfp57*, and in which expression of *Fgfr2* is more variable than in the EU cells, but do not appear to have specified either PrE or Epi lineages. We propose that this cluster represents cells in the process of <u>m</u>aking a <u>d</u>ecision (MD).

At the early blastocyst stage the majority of wild-type cells are in Cluster 5 (MD), with only 27% of cells in Cluster 2 (EU; Fig. 3C). In contrast the majority (72%) of null cells from early blastocysts are in the EU Cluster, with only 18% in the decision-making stage. In later stage blastocysts the majority of wild-type cells have committed to either primitive endoderm or epiblast, while 29% are in the decision making phase. The majority of *Sall4*-null cells in later blastocysts have left the EU category and begun the decision-making process, however few of them have successfully left this state and managed to specify either lineage (Fig. 3C). Notably, those *Sall4*-null cells located in the PrE or Epi clusters do not segregate away from the wild type cells based upon their gene expression patterns, indicating that *Sall4*-null blastomeres are capable of successfully specifying either PrE or Epi lineages (Fig. 3B). This is consistent with our observation that some cells in implanting blastocysts are able to express multiple markers of primitive endoderm or epiblast and occupy an appropriate spatial position within the embryo (Fig. 2B, S1F). Yet the reduced proportion of *Sall4*-null cells in either of these two categories indicates that their efficiency of making this decision is reduced.

While wild type and *Sall4*-null cells do not notably segregate by genotype in most of the cell state categories in Figure 3B, cells in the EU stage are almost completely segregated by genotype. This indicates that Sall4 has a particularly pronounced influence on gene expression patterns at this stage. Cells about to undergo a fate transition have been found to show increased transcriptional noise or a decrease in cell-cell correlation (Mohammed et al., 2017; Mojtahedi et al., 2016; Richard et al., 2016). While we do find a dip in cell-cell correlation at the EU stage, *Sall4*-null EU cells show a significantly higher average correlation than do WT cells, which is consistent with null cells being less likely to undergo a fate transition (Fig. 3D). Curiously, *Sall4*-null Epi cells also show a significant increase in cell-cell correlation as compared to wild type cells, which could explain the decreased efficiency of ES cell derivation from Sall4-null epiblasts (AM and BH unpublished and (Tsubooka et al., 2009)). Together these data indicate Sall4 plays an important function in stabilising gene expression patterns in unspecified ICM cells (the EU state).

The nature of the gene expression defect can be visualised by comparing gene expression patterns in cells based upon developmental stage (i.e. cluster; Fig. 4A). Most genes assayed show a significant decrease in gene expression in *Sall4*-null EU cells, whereas those cells that have achieved the PrE or Epi stages show normal expression patterns (Fig. 4A). Cells in the process of making the cell fate decision (MD) show a near-normal expression pattern for most genes assayed, but nevertheless still show reduced expression of important EPI (*Esrrb, Sox2, Klf2*) and PrE (*Gata4, Sox17*) markers. We therefore conclude that Sall4 activity is not required for cells to adopt a primitive endoderm or epiblast fate per se, but rather is required to establish appropriate gene expression patterns in early ICM cells to maximise the efficiency of the PrE vs Epi decision-making process (Fig. 4B).

**Figure 4.**
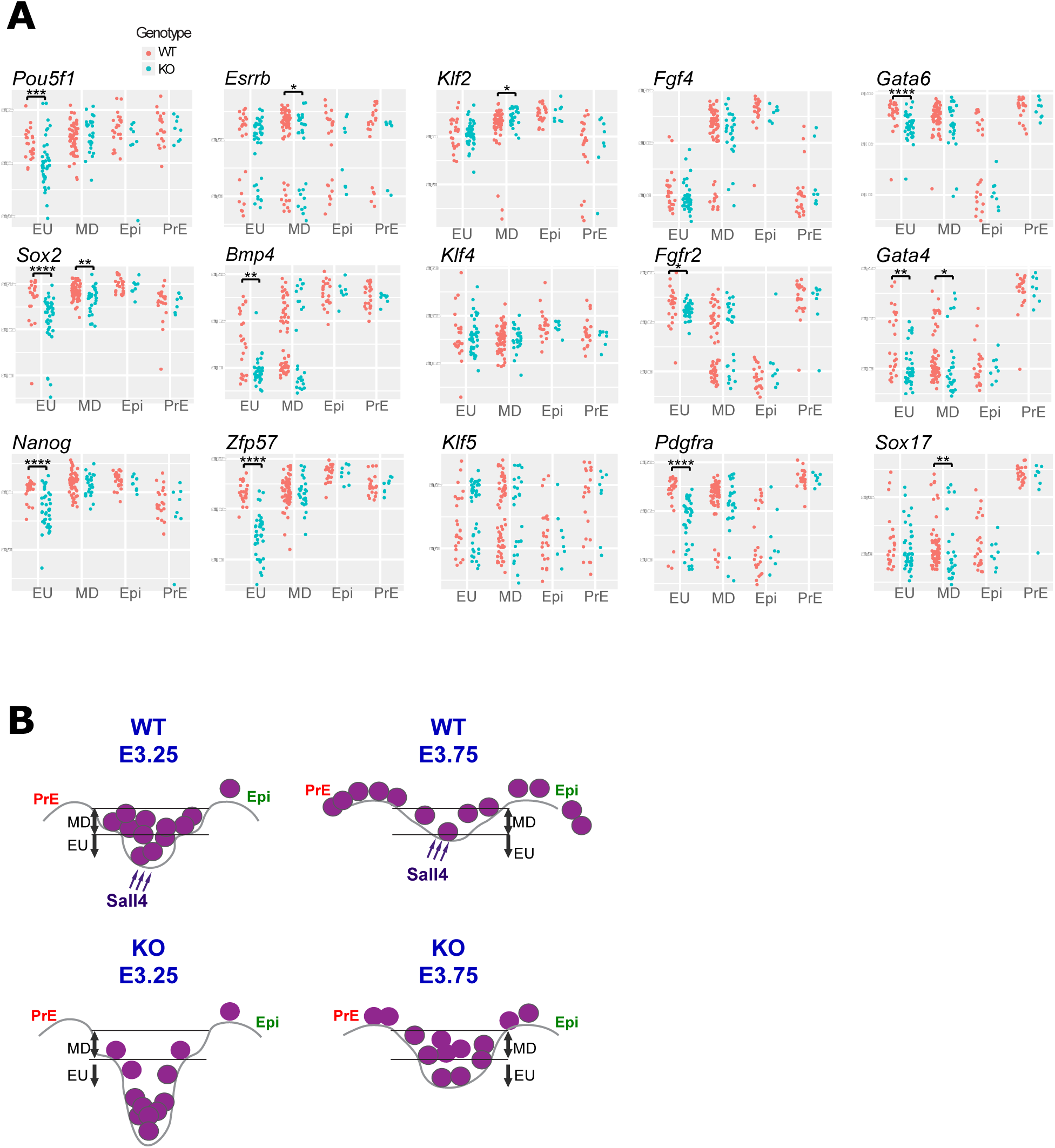
Sall4 promotes gene expression in early unspecified ICM cells. (A)Single cell expression data for genes characteristic of Epiblast and Primitive Endoderm split into clusters as specified in Fig. 3B for control (WT, red) and Sall4-null (KO, Blue) cells. Each dot represents a single cell. Statistical significance was calculated using a Kruskall-Wallis test (Supplemental Table 1): * P ≤0.05, ** P ≤0.01, *** P ≤0.001, **** P <=0.0001. (B)Model of how Sall4 facilitates lineage commitment. In early wild type ICMs (top left) most cells (purple circles) exist in an attractor state of a differentiation landscape associated with an unspecified identity (EU), though some cells occupy the MD state, which is essentially a ‘staging post’ en route towards differentiation. The probability of achieving a differentiated state (PrE or Epi) is very low. As the embryo matures (top right) the depth of the attractor state lessens and cells in the MD state are able to commit to either the PrE or Epi state. In the absence of Sall4 (bottom panels) the depth of the EU attractor state is more pronounced, and cells within this state show increased cell-cell variability. As the embryo matures the depth of the attractor is lessened, but remains deeper than in wild type cells so most Sall4-null cells still occupy the MD state and few have committed to either PrE or Epi. Black horizontal lines represent the threshold limits of the EU and MD states.

We propose that the perturbed GRN observed in *Sall4*-null EU cells results in them existing in an abnormally stable attractor state, visualised in Fig. 4B as an increase in the depth of the energy well in which they exist in a differentiation landscape. While both wild type and *Sall4*-null cells respond to signalling cues associated with ICM maturation and the Epi vs PrE lineage commitment step (i.e. FGF signalling), these differentiation cues are insufficient to drive cells out of the deeper well in the *Sall4*-null state, through the MD intermediate and into a new cell state (Fig. 4B). At low frequency some *Sall4*-null cells are able to escape this early developmental state whereupon they can achieve the PrE or Epi attractor state and establish a near-normal GRN. Sall4 is therefore not required for cells to activate the EPI or PrE GRNs specifically, but rather Sall4 facilitates the exit out of the unspecified state (Fig. 4B). This is consistent with the observation that *Sall4*-null ICMs are competent for ES cell derivation, although at reduced frequency and efficiency (AM and BH, unpublished observations and (Tsubooka et al 2009)). Once established, *Sall4*-null naïve ES cells are able to self-renew and to maintain the pluripotency GRN, though again require Sall4 activity when exiting the self-renewing state (Miller et al., 2016).

Establishing correct gene expression patterns at the appropriate time underlies successful cell fate transitions, which are the basis of developmental progression. We show here that Sall4 is required to maintain appropriate expression levels of multiple genes in the unspecified progenitor pool of ICM cells in early blastocysts. In its absence fewer cells are able to make the epiblast vs primitive endoderm decision, resulting in a skewing of tissue distribution in the implanting blastocyst. This demonstrates the importance of setting up correct gene expression patterns early in development in order for cells to make the correct decisions at the correct time.

---

**Figure S1.**
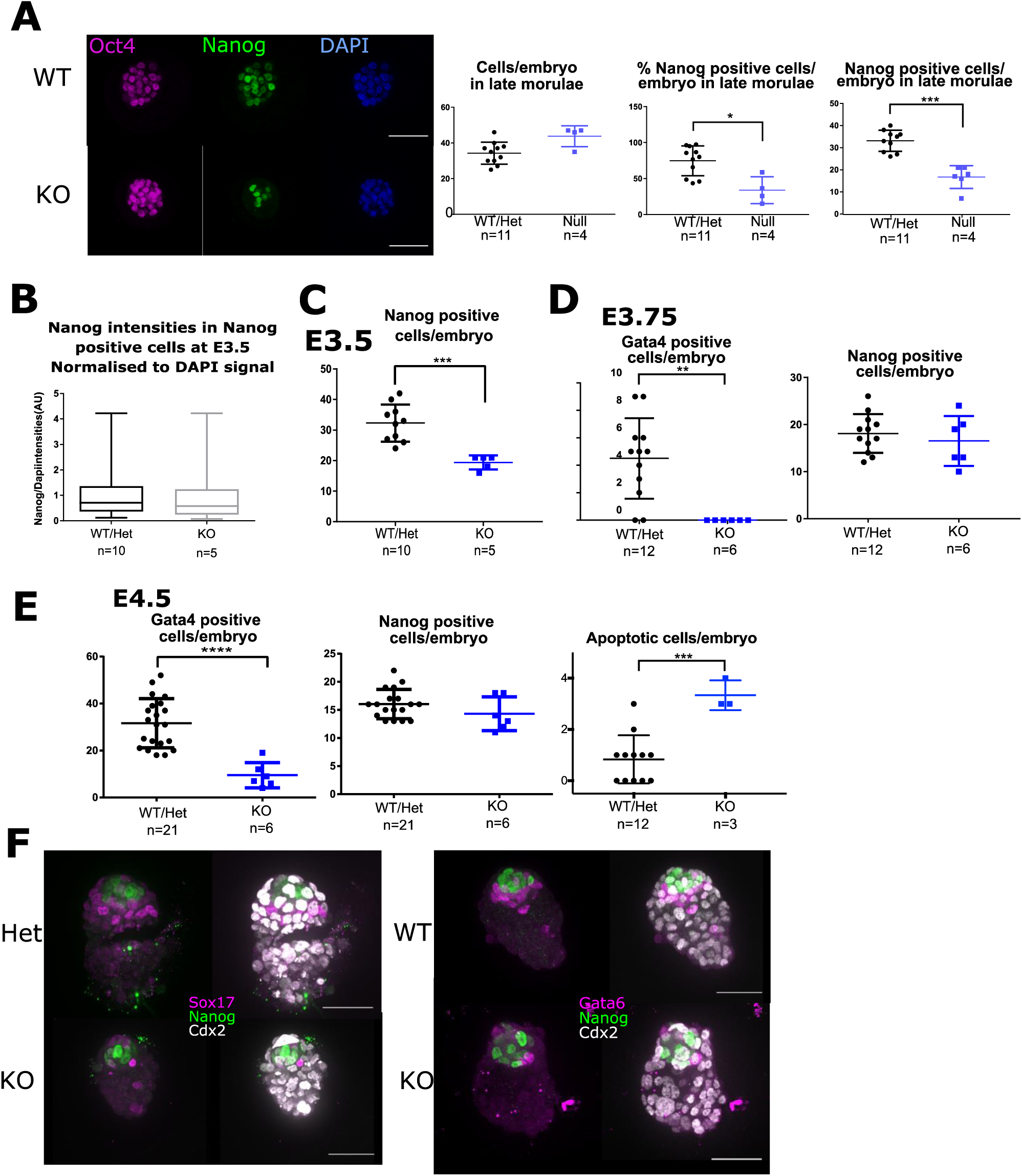
(A)(Left) Representative projection images of anti-Oct4 (magenta), anti-Nanog (green) and DAPI (blue) staining in control embryos (WT/Het n=11) and Sall4^Δ/Δ^ (KO, n=4) late morula embryos. Scale bar = 50μM. (Right) Quantification of the embryos showing number of cells per embryo, the percentage of Nanog positive cells per embryo and the absolute number of Nanog positive cells per embryo. Significance was determined using a Mann Whitney test for all three comparisons (* P≤0.05, *** P≤ 0.001). (B)The intensity values for the Nanog positive cells from (Fig. 1B) normalised to DAPI signal for control (Wt/Het) and KO embryos. (C)Quantification of Fig. 1B showing absolute number of Nanog positive cells per embryo. Significance was determined using a Mann Whitney test for all three comparisons (*** P≤ 0.001). (D)Quantification of Fig. 1D showing absolute number of Gata4 positive cells (left) and Nanog positive cells (right) per embryo. Significance was determined using a Mann Whitney test for all three comparisons (** P≤ 0.01 (E)Quantification showing absolute number of Gata4 positive cells (left) and Nanog positive cells (middle) and Apoptotic cells (as judged by Caspase3 staining; right) in 4.5 dpc embryos. Significance was determined using a Mann Whitney test for all three comparisons (*** P≤0.001, **** P≤ 0.0001). (F)(Left) Representative projection images of anti-Nanog (green), anti-Sox17 (magenta) and anti-Cdx2 (white) staining in control (WT/Het n=4) and Sall4^Δ/Δ^ (KO, n=4) E4.5 late blastocysts. (Right) Representative projection images of anti-Nanog (green), anti-Gata6 (magenta) and anti-Cdx2 (white) staining in control (WT/Het n=27) and Sall4^Δ/Δ^ (KO, n=6) E4.5 late blastocysts. Scale bar = 50μM.

**Figure S2.**
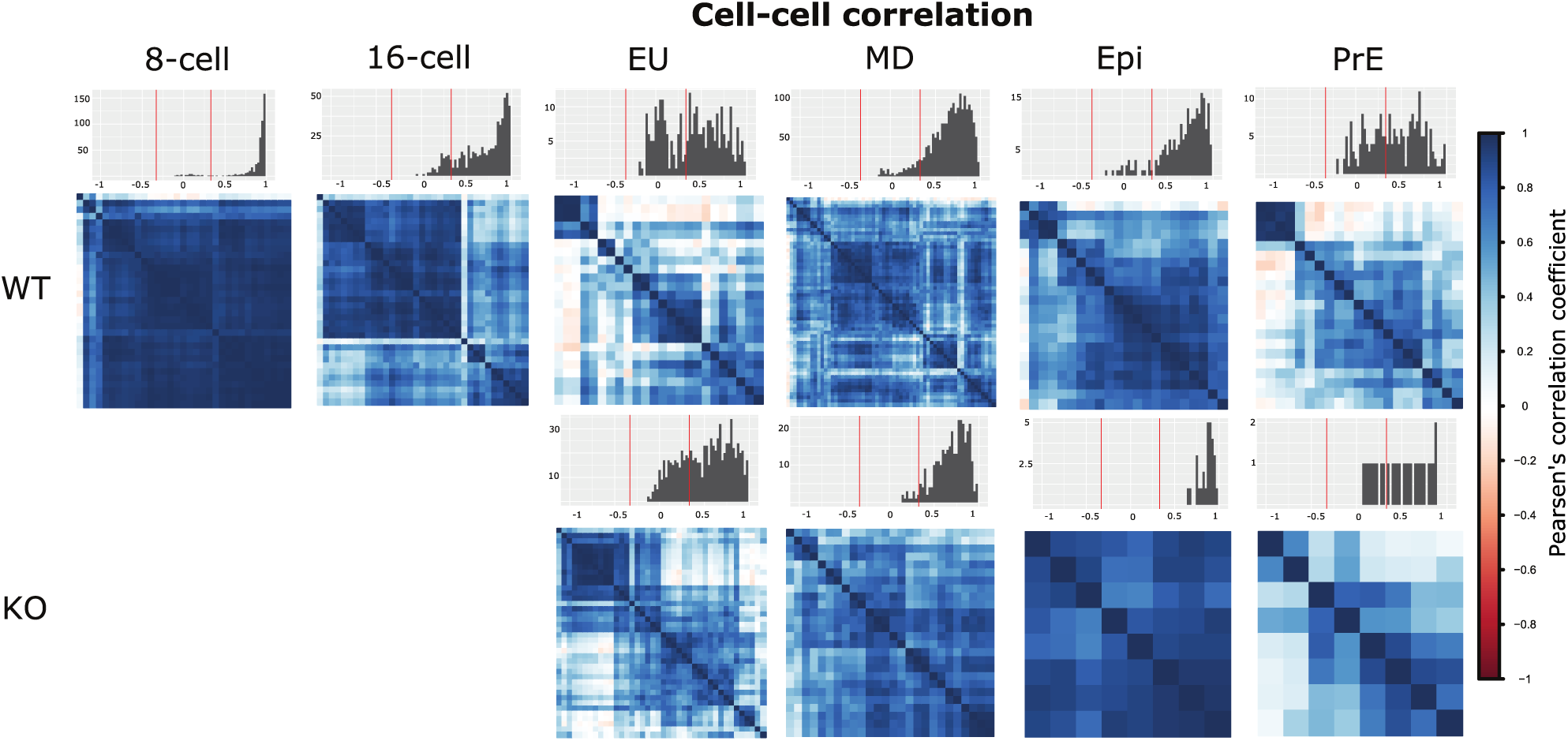
Cell-cell correlation heatmap. Pearsen’s correlation coefficients of each cell are compared to each cell in each Category (EU, MD, Epi, PrE) for KO and WT cells. Above each correlation matrix is a histogram displaying the distribution of correlation coefficients. The red lines on the histograms are to illustrate anticorrelated (-1 to -0.33), no correlation (-0.33 to 0.33) and positive correlation (0.33 to 1).

**Supplemental Table 1:** Result of Kruskall-Wallis tests comparing gene expression at different time points and developmental stages.

**Supplemental Table 2:** Raw CT values for single cell gene expression data used in this study.

## Materials and Methods

### Mice and embryos

All animal experiments were approved by the Animal Welfare and Ethical Review Body of the University of Cambridge and carried out under appropriate UK Home Office licenses. *Sall4^Flox/Flox^* mice (Sakaki-Yumoto et al., 2006) were crossed to a line of mice expressing Sox2-Cre (Hayashi et al., 2002) to delete the Floxed allele, creating *Sall4^Δ/+^* mice which were then maintained by crossing to the C57Bl/6 strain (backcrossed for at least 3 generations). These mice were genotyped using DNA extracted from ear biopsies using a duplex PCR reaction with the following primers: 5’-CCTCCCGGAATTGCTTATCT-3’, 5’-GGAGAGAAGCCTTTCGTGTG-3’, 5’-TGGTCTACAGTGCAAGTGCC-3’. Embryos were generated by natural mating, where successful mating was confirmed by the detection of a copulation plug. Embryo staging was based on the assumption that mating occurred at midnight so that at midday the embryos were assigned embryonic day (E) 0.5. Embryos were collected at the relevant stages from the oviduct or uterus in M2 medium (Sigma). The zona pellucida was removed if necessary using Acid Tyrode’s solution. To genotype the embryos a duplex nested PCR was formed, with the following primers used for the first PCRrun:5’-AACATGGGACAGCAGTGAATG-3’,5’-GCAGATCCACGAGCGAACAC-3’,5’-TCACACAAAGCAAGACTTTTACCGCT-3’. The second PCR run uses the same primers as to genotype the mice stated above.

To culture embryos in exogenous Fgf, embryos were flushed from plugged super-ovulated females (5-6 weeks old using PMSG/hCG) at the 8-cell stage (E2.5), and cultured in M16 media with or without the presence of 500 ng/μl or 1 μg/μl of FGF2 (made in house). They were incubated at 37°C, 5% C02 for 48 hours.

### Immunofluorescence

Embryos were fixed in 2.5% paraformaldehyde, permeabilised in 0.25% Triton X-100 and 3% polyvinlypyrrolidone in PBS, and blocked in PBS containing 10% fetal bovine serum and 0.1% Triton X-100. The embryos were incubated in primary antibodies in blocking solution at 4°C overnight, and secondary antibodies for an hour at room temperature. Primary antibodies were used at the following dilutions: anti-Oct4 (1/200, Santa Cruz Biotechnologies, sc-8628), anti-Nanog (1/200, Cosmo Bio, RCAB002P), anti-Cdx2 (1/200, Abcam, ab157524), anti-Gata6 (1/200, R&D Systems, AF1700), anti-Gata4 (1/200, Santa Cruz, sc-1237), anti-Sox17 (1/200, R&D Systems, AF1924), anti-Sall4 (1/200, Cosmo Bio, PPX-PP-PPZ0601-00), anti-cleaved caspase 3 (1/500, Cell Signaling Technologies, 9664S).

### Imaging and image analysis

Embryos were imaged in Ibidi μ-slides (Thistle Scientific). Images were taken on the Spinning Disc (Andor Revolution XD microscope) or the confocal (Leica SP5 confocal microscope). Intensity values for each cell in the embryo, and total cell numbers were analysed using MINS software (Lou et al., 2014). Other cell counts were performed manually in ImageJ. Plots were generated in Graphpad Prism, and statistical significance was tested using an unpaired Mann-Whitney test. Sall4 heterozygous embryos showed no early embryonic phenotype, and as such for all experiments they were grouped with WT embryos.

### Single cell analysis

Embryos were flushed with M2 media from plugged super-ovulated females (5-6 weeks old using PMSG/hCG) either at E3.25 (6am on Day 4 post coitus) or E3.75 (6pm on Day 4 post coitus). The zona pellucida was removed using Acid Tyrode’s solution, and immunosurgery performed to remove the trophectoderm from the ICM. The trophectoderm lysate was used to genotype the embryos.Each ICM was dissociated into single cells using a drop of 1:1 mix of 0.025% trypsin-EDTA and 0.025% trypsin-1% chick serum overlaid with mineral oil, incubated for 10 minutes at 37°C, 5% CO2. Partially dissociated cells were washed in N2B27 + 10μM Hepes + 1.5mg/ml BSA and single cells mechanically dissociated using finely drawn glass capillaries. Pre-amplication, cDNA synthesis and real-time PCR was performed as described in (O’Shaughnessy-Kirwan et al., 2015). Three housekeeping genes were used for normalisation (*Ppia, Hprt* and *Gapdh*). Only cells that showed expression of all these genes were used for subsequent analysis. Raw Ct values are provided in Supplementary Table 2, and the Taqman probes used in this study are as described in (O’Shaughnessy-Kirwan et al., 2015). N= 44 KO cells from 8 embryos, 62 WT/Het cells from 10 embryos. For Data in Figure 4A, KO EU cells N=40 (from 7 embryos), KO MD N=26 (7 embryos), KO Epi N=8 (4 embryos), KO PrE N=8 (3 embryos), WT EU N=24 (13 embryos), WT MD N=61 (20 embryos), WT Epi N=22 (11 embryos), WT PrE N=21 (8 embryos).

All data manipulations and analyses were performed using R. An empirically defined limit of detection was set at Ct=28. The mean of the housekeeping genes was subtracted from each sample to create dCt values, which were transformed using 2^-dCt^. Hierarchical clustering was performed using ‘pheatmap’ package, and diffusion maps were generated using ‘destiny’ package. The cell-cell correlation analysis used Pearsen’s correlation coefficient using the “corrplot” package, using only the absolute value for Fig3D. All other plots were generated using ‘ggplot2’ package.

#### Acknowledgements

We thank Peter Humphreys and SCI Tissue Culture Staff for excellent technical assistance; members of the B.H. lab for stimulating discussions; and Alfonso Martinez-Arias, Carla Mulas, Jenny Nichols, Berenika Plusa, Nicola Reynolds, Veronica Biga and Austin Smith for critical comments on the manuscript.

## Author Contributions

A.M. and B.H. conceived the project, A.M., S.G., C.E.-D. and B.H. performed the experiments; A.M. performed all quantitative analyses; RN provided reagents and A.M. and B.H. wrote the paper.

